# GEMBS – high through-put processing for DNA methylation data from Whole Genome Bisulfite Sequencing (WGBS)

**DOI:** 10.1101/201988

**Authors:** Angelika Merkel, Marcos Fernández-Callejo, Eloi Casals, Santiago Marco-Sola, Ronald Schuyler, Ivo G. Gut, Simon Heath

## INTRODUCTION

DNA methylation is essential for normal embryogenesis and development in mammals and most commonly found at CpG dinucleotides. Whole genome sequencing of bisulfite converted DNA (WGBS) currently represents the gold standard for studying DNA methylation at genomic level as contrary to other techniques, it provides an unbiased view of the entire genome at single base pair resolution (see Bock *et al.* 2012). In practice, due to its comparatively high cost, its application for the analysis of large data sets (i.e. > 50 samples) has been lagging behind other more cost-efficient platforms, such as for example methylation microarrays (e.g. Infinium 27K, 450k and EPIC). Despite the variety of software tools that exist for the analysis of WGBS, processing of large datasets is cumbersome hampered by inefficiency as well as limited functionality of the available tools (see Krueger *et al,* 2012 for a review on WGBS analysis tools). As a result of decreasing costs for sequencing and computational infrastructure, the frequent incorporation of WGBS as a standard assay in large epigenomic consortia such as BLUEPRINT, ENCODE or NIH ROADMAP is now calling for the development of a new generation of analysis tools.

Here, we present GEMBS, a bioinformatics pipeline designed for the analysis of large WGBS data sets with specific focus on computational performance and the implementation of common analysis standards. It combines two functionalities: 1) a high performance read aligner called GEM3 (Marco-Sola et al. 2012, Marco-Sola 2017), and 2) a variant caller specifically developed for bisulfite sequencing data named BScall. Both components are embedded in a state-of-the-art, highly efficient and parallelizable work flow that ensures fast and reliable execution. We show that GEMBS greatly outperforms other currently available tools and demonstrate how GEMBS can be used for accurate variant calling from WGBS data.

Analysis of bisulfite sequencing data starts with the alignment of bisulfite converted reads, and although powerful tools for short read alignment exists (e.g. Bowtie, BWA, SOAP, etc.), the peculiarities of bisulfite converted sequences, such as increased rate of mismatches and low sequence complexity, prevent a straightforward implementation. Bisulfite treatment converts un-methylated cytosines to uracils in the original DNA strands, which are replaced by thymines during PCR amplification. This creates four potentially different sequences from a single stretch of DNA that need to be aligned to the reference genome sequence. Directional libraries ensure the selective sequencing of the original strands (rather than the amplicons) via adaptor tagging. However, using a conventional short read aligner for mapping, the converted bases would be interpreted as mismatches, and the large percentage of these (since most cytosine are unmethylated typically approaching 25% of the sequence) would result in a large number of unmapped or poorly mapped reads.

Two solutions have been proposed to handle the mapping problem. So called ‘3-letter’ aligners, such as Bismark (Krueger et al., 2011), BWA-meth (Pedersen et al., 2014) and Novoalign (www.novoalign.com), perform a two stage mapping process: Firstly, C depleted reads (from read 1) are ‘fully converted’ by converting all remaining C’s to T’s before mapping, while G depleted reads (from read 2) have all remaining G’s converted to A’s. Secondly, using a classic short read aligner, mapping is then performed to two altered versions of the desired reference sequence, one with all C’s converted to T’s, and the other with all G’s converted to A’s.^1^ After mapping, the original sequence data should then be restored before downstream analysis. Alternatively, ‘methylation aware’ aligners or ‘4-letter’ aligners, such as BSMAP (Xi and Li, 2009), Last (Frith et al., 2012), and GSNAP (Wu and Nacu, 2010), consider both cytosines and thymines as potential matches e.g. by creating multiple seeds during indexing (BSMAP; Xi and Li, 2009). Some tools besides employ multiple alignment options to accommodate advantages and disadvantages of each approach and to additionally enable the analysis of color space data, e.g. BSmooth (Hansen et al 2012) or MethylCoder (Pedersen et al 2011) (see Krueger et al, 2012).

GEMBS implements the first approach in a straight forward efficient manner. All of the conversion steps before and after mapping are performed on the fly on a read-pair-by-read-pair basis in the mapper GEM3 itself. Additionally, GEM3 only performs one alignment against a single composite reference. Its internal design allows GEM3 to handle large indices, where other aligners cannot and hence require two alignments: one against the converted and one against the unconverted reference. Both strategies avoid generation of intermediate files and reduce unnecessary processing steps.

After successful read alignment, the next analysis step is the determination of cytosine methylation status. It is commonly determined as the ratio of reads with an unconverted cytosine (i.e. C) over the sum of all reads containing either an unconverted cytosine or a converted cytosine (i.e. T). Aside from confounding genetic variants (and mapping errors), misinterpretations may arise from base calling errors and over- or under-conversion of cytosines during the bisulfite treatment (that is methylated cytosines which should be resistant to the conversion are converted and susceptible un-methylated cytosines which should be converted are not converted, respectively).

One possibility is to filter for known SNPs either derived from sample matched genome sequencing data (e.g. Stadler *et al,* 2012), SNP arrays or public databases such as dbSNP (https://www.ncbi.nlm.nih.gov/projects/SNP/). Alternatively, genotypes can be estimated from the same WGBS data that is used for calculating the methylation proportion, e.g. Bis-SNP (Liu *et al,* 2012), MethylExtract (Barturen *et al,* 2013) or BS-SNPer (Gao *et al,* 2015).

With BScall, we have adopted a Bayesian model similar to that used in Bis-SNP to model the conversion process (see METHODS). We use this model to simultaneously infer the most likely genotype and most likely methylation proportion given base error probabilities, and under/over conversion rates. In the interest of computational performance, our current BScall implementation does not perform local realignment as it is implemented for example Bis-SNP. Although realignment increases sensitivity for indel detection our focus lies on true cytosines not variants, hence we accept faulty SNP detection to a certain degree. Further, BScall does not incorporate prior genotype probabilities as they may lead to non-replicable results (i.e. when SNP frequency are taken from public data bases that undergo periodic updates, e.g. Bis-SNP) and in any case may be incorporated post-analysis.

## RESULTS

### GEMBS workflow and implementation

Mapper (GEM3) and caller (BScall) are both implemented in the GEMBS analysis pipeline. The pipeline automates data processing efficiently and reliably employing state-of-the-art tools and data standards for HTS analysis, e.g. Samtools, BCFtools (Figure 1).

**Figure 1.**
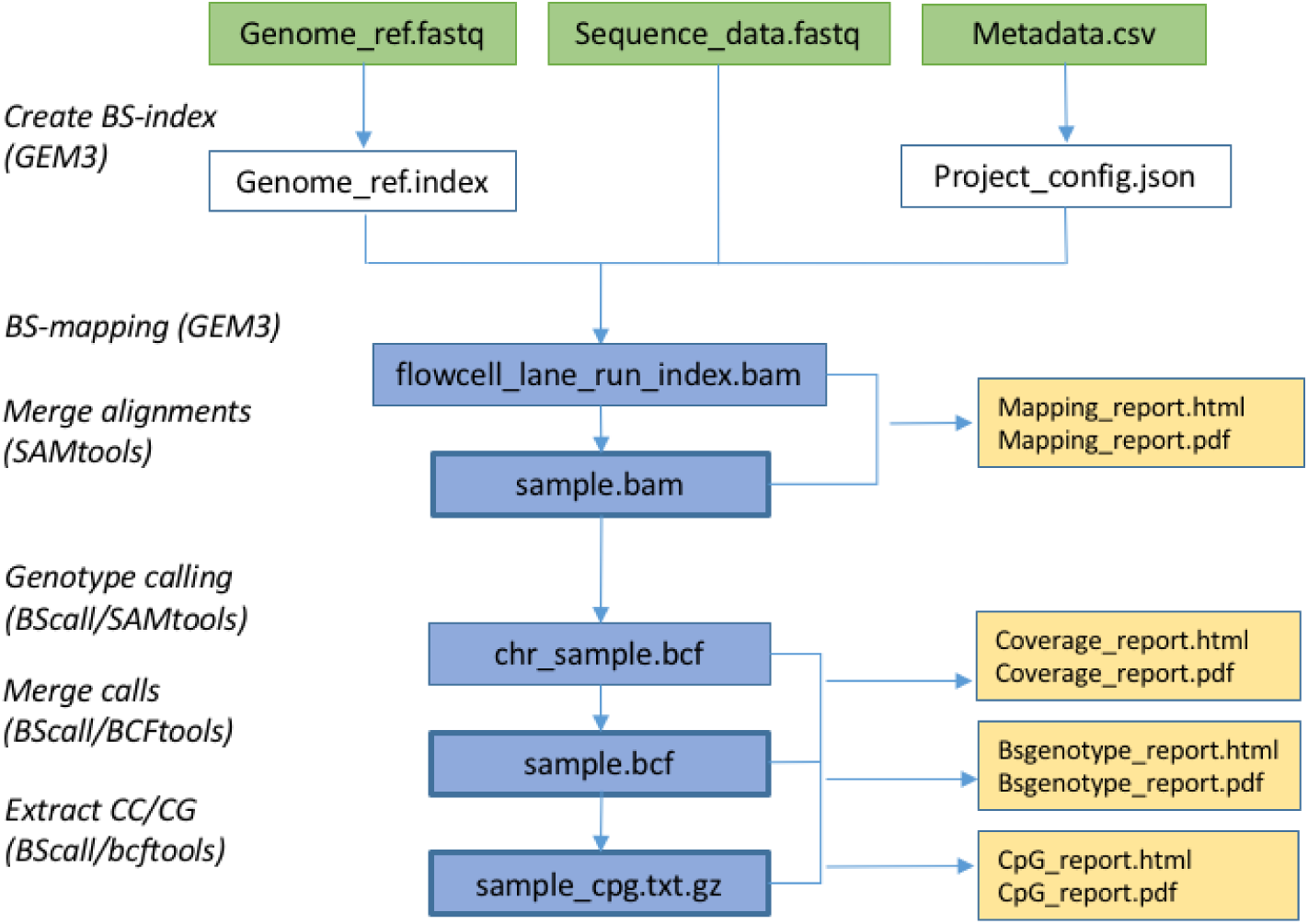
Overview of the GEMBS workflow. Processes and tools in italic letters. Files in colored boxes (green = input; white = pre-processing, blue = output; yellow = quality control)

GEMBS requires as input 1) a genome reference, 2) meta data describing the project, i.e. the layout of the sequence data (multiplex index, library, lanes per samples, samples per project) and 3) the sequence data (Figure 1A). In a pre-processing step, a bisulfite index of the reference is built for fast and efficient read mapping, and the meta data together with a set of preset parameters is used to construct a configuration file from which all software components are run.

Reads are mapped with GEM3 in bisulfite mode and the output is concatenated into standard alignment files (.bam format) according to the sample description (e.g. all lanes per chromosomes or all chromosomes per sample). In order to compensate for the low sequence complexity, we allow for as much as 10% mismatches. Unwanted adaptor sequences and low quality base pairs are excluded by soft trimming of the 5’ and/or 3’ read ends, respectively (benchmarking shows hard trimming is more aggressive and of lower performance). From the alignments, BScall produces genotype calls and stores them in .bcf format alongside with strand-specific information for each base call (homozygous AA and TT calls are not reported is they are not relevant for BS-seq analysis)^2^. Only uniquely mapping reads (mappability score >= Q20) are used for the analysis (in the case of pair end sequencing only uniquely mapping pairs in the correct orientation and on the same chromosome). Duplicates with identical start and end coordinates are collapsed into singletons and the first 5 nucleotides are removed from the 5′ end of each read to eliminate artefacts from library preparation (end repair). Ultimately, all CC/GG dinucleotide positions are extracted and stored in a separate text file.

For each of the pipeline outputs (mapping, genotypes, CpG) a set of QC statistics is calculated, stored initially as .json format and later published as .pdf of .html report. Instead of performing a separate QC analysis after the composition of the output as it is done in most analysis tools, collection of the QC statistics happens in parallel with the mapping and calling process at no computational overhead. As QC metrics, we adapted standard QC measures recommend by GATK and the IHEC consortium.

Genotype calls (‘SNP.bcf’) and CpG estimates (‘CpG.txt.gz’) serve for subsequent downstream analysis. Cytosine methylation in non-CpG context can be easily derived from the genotype calls, similarly allele specific methylation. The reported CpGs are usually further filtered according to user preferences, mostly by minimum read coverage. We have developed additional software for methylation analyses, such as differential methylation analysis and methylome segmentation, as well as secondary processing and functional annotation which can be made available upon request. Alternatively, GEMBS output can be easily converted for analysis with other popular tools, such as MethylKit, Bumphunter, MethylSeekR, etc. (Akalin *et al* 2012, Jaffe et al 2012, Burger *et al,* 2013). Finally, GEMBS creates standard bigWig files for visualization of CpG methylation, methylation standard deviation and coverage that can be displayed as custom tracks in the UCSC genome browser (https://genome.ucsc.edu/).

Although it is possible to run GEMBS steps sequentially rather than in parallel when only limited resources are available, we recommend at minimum 32GB RAM and 1 TB of disc space for processing a standard 30X human WGBS dataset. GEMBS achieves best performance results when deployed on a distributed computer system (computer cluster) not only because multiple workflows can be executed in parallel (e.g. multiple chromosomes, samples) but because the software components internally employ parallelization and multithreading. For the analysis of large data sets, execution of GEMBS can be further automated. For example, for libraries sequenced *in house* we import project meta data directly via API from our Laboratory Information Management System (LIMS). Similarly, meta data might be imported from any data server given the appropriate format. Secondly, we employ the *‘JIP pipeline system’* (http://pyjip.readthedocs.io/en/latest/), a pipeline management system to manage the large number of workflows on the computer cluster and to ensure efficient and trackable analysis. In principle any pipeline management system can be implemented as long as it is adapted to the individual cluster architecture (for an alternative check https://www.nextflow.io/). Using our implementation, we have achieved processing of, for example, 36 WGBS datasets at 30X coverage in less than 3 days.

### Performance

To evaluate the performance of GEMBS, we separately benchmarked our aligner GEM3 and caller BScall against some of the popular tools for WGBS data analysis (see Table 1).

**Table 1.** Tools overview

| Task | Software | Version | Reference |
| --- | --- | --- | --- |
| Read alignment | Bismark | Bismark 0.16.1 | Krueger et al. (2011) |
|  |  | bowtie2 2.2.9 | Langmead et al (2012) |
|  | BSMAP | BSMAP 2.90 | Xi et al. (2009) |
|  | Novoalign | Novoalign 3.05.01 | http://www.novocraft.com/ |
|  | BWAmeth | BWAmeth 0.10 | Pedersen (2014), Li (2013) |
|  |  | bwa 0.7.7-r441* |  |
|  | GemBS | GemBS 1.2.2 |  |
|  |  | GEM3 v3.1.0 | Marco-Sola et al. (2013), Marco-Sola (2017) |
| SNP and methylation calling | Bis-SNP | BisSNP 0.82.2 | Liu et al. (2012) |
|  | MethylExtract | MethylExtract 1.8.2 | Barturen et al (2013) |

We started by comparing the BS-alignments (Figure 2–3, SupplTable S1, S2) and found that all mappers were able to align at least 80% of the sequenced bases and that sequencing coverage had no influence on sensitivity or specificity of the alignments. GEM3 (GEMBS) emerges as the fastest mapper aligning 85% of all sequenced bases with a high alignment quality score (MapQ>20) in less than three hours. BSMAP and Novoalign are similarly sensitive but slower than GEM3 (e.g. CPU time for Novoalign of 100 and 200 days for 27X and 58X, respectively). Bwa-mem and Bismark rank third and fourth in speed but last in the alignments^3^. This might lead to the conclusion that when sensitivity is required slower processing could be accepted. Note here, however, that the definition of uniquely mapped reads is not uniform across mappers and can be misleading when for example only a fraction of the mappings is reported.

**Figure 2.**
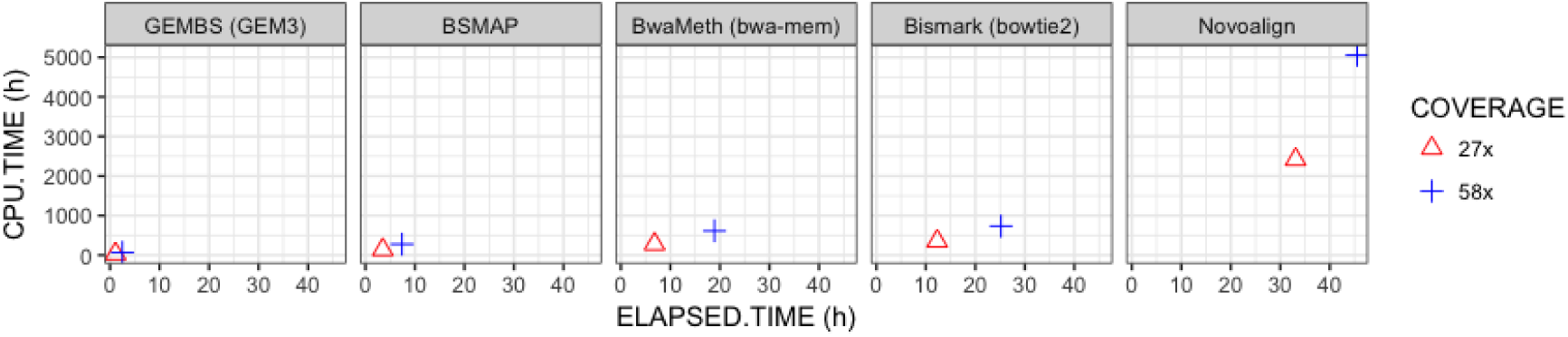
Alignment times per mapper for 27X and 58X coverage

**Figure 3.**
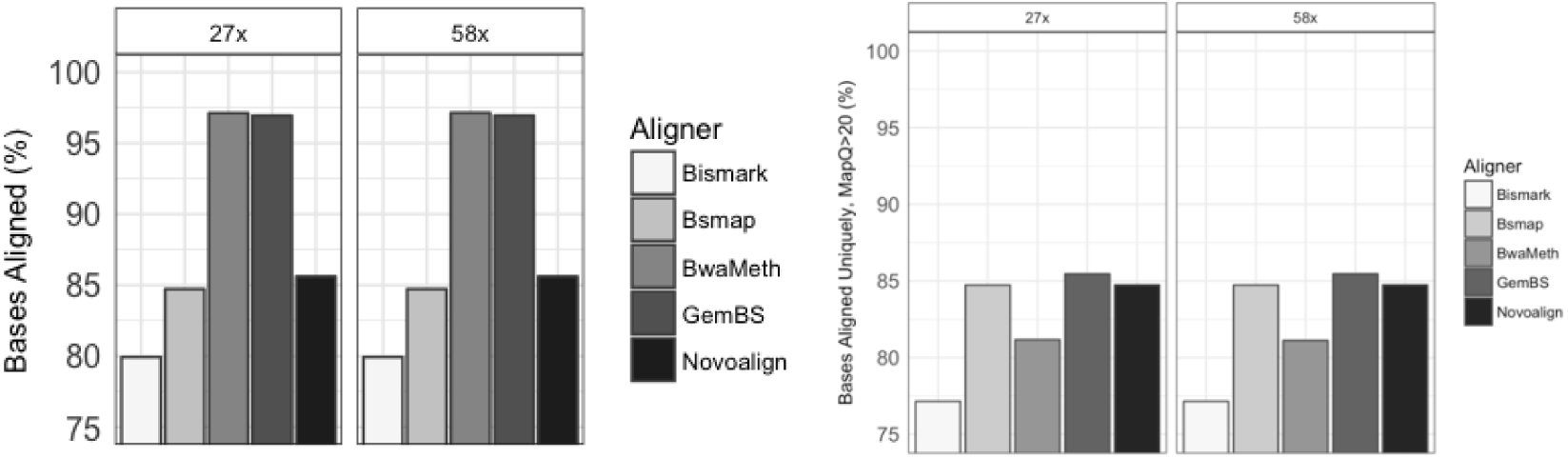
Read alignments per mapper for 27X and 58X coverage. Left: Bases aligned, Right: Uniquely mapped bases (mapQ>20)

Continuing the benchmark across callers, we find the differences in processing times are far greater than those between mappers (Figure 4, SupplTable 3). BScall processes the 58X data set in less than 3 hours whereas the Bis-SNP and MethylExtract require several days independent of the mapper that produced the alignment. Similarly, the number of SNPs detected depends in first instance on the caller used, then the underlying alignments and sequencing coverage. The large number of SNPs produced by BScall (followed by MethylExtract then Bis-SNP), especially in combination with BSMAP alignments are most likely a result of a large number of false positive (Figure 5). Without local realignment implemented, BScall may misinterpret indels. BSMAP produces a large number of multi mapping reads by alignment against thymines and cytosines whereby incorrect MapQ calibration easily leads to false SNP calls. Quality filtering hence becomes crucial. For this comparison, we have imposed only default parameters (see METHODs) resulting in strict filtering for Bis-SNP, but none or very little for GEMBS and MethylExtract, respectively.

**Figure 4.**
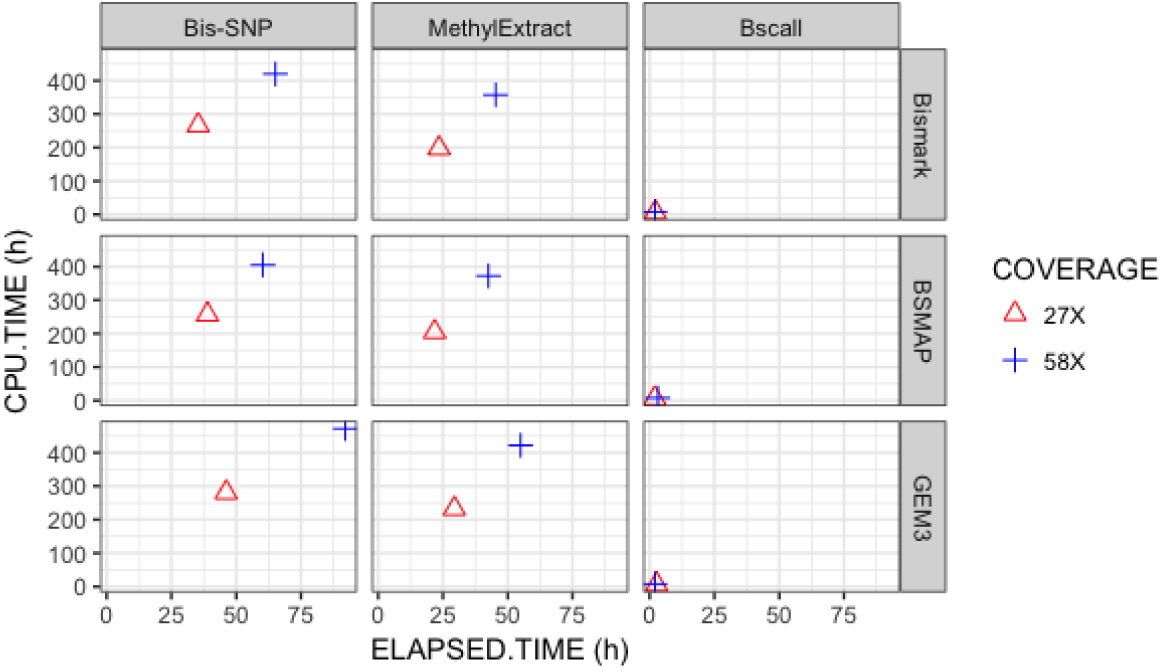
Processing times for SNP calls across different aligners and callers

**Figure 4.**
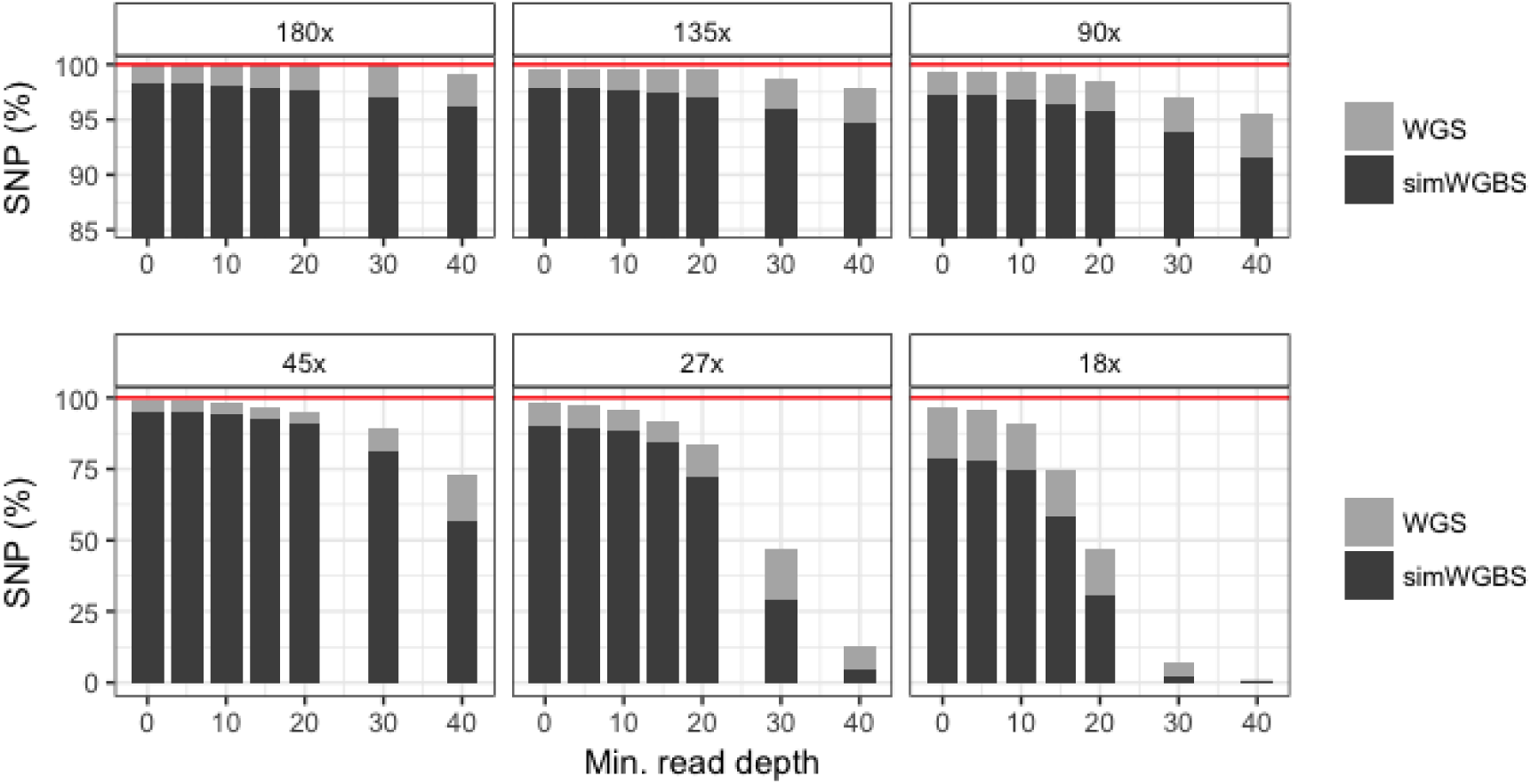
Number of SNPs called by BScall (WGBS) and FreeBayes (WGS) at multiple sequencing depth

**Figure 5.**
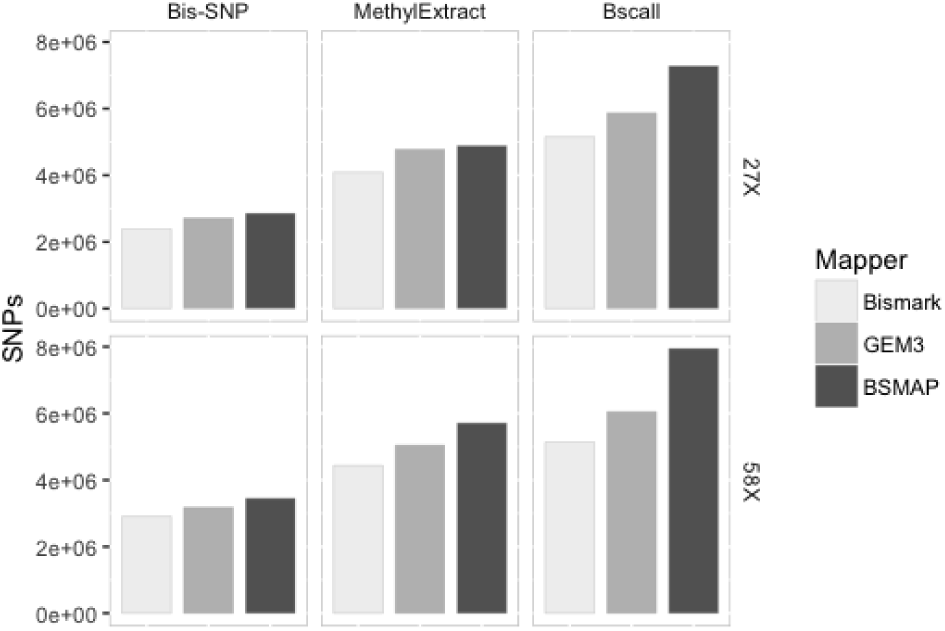
SNP calls by mapper/caller for 27X and 58X coverage

**Figure 5.**
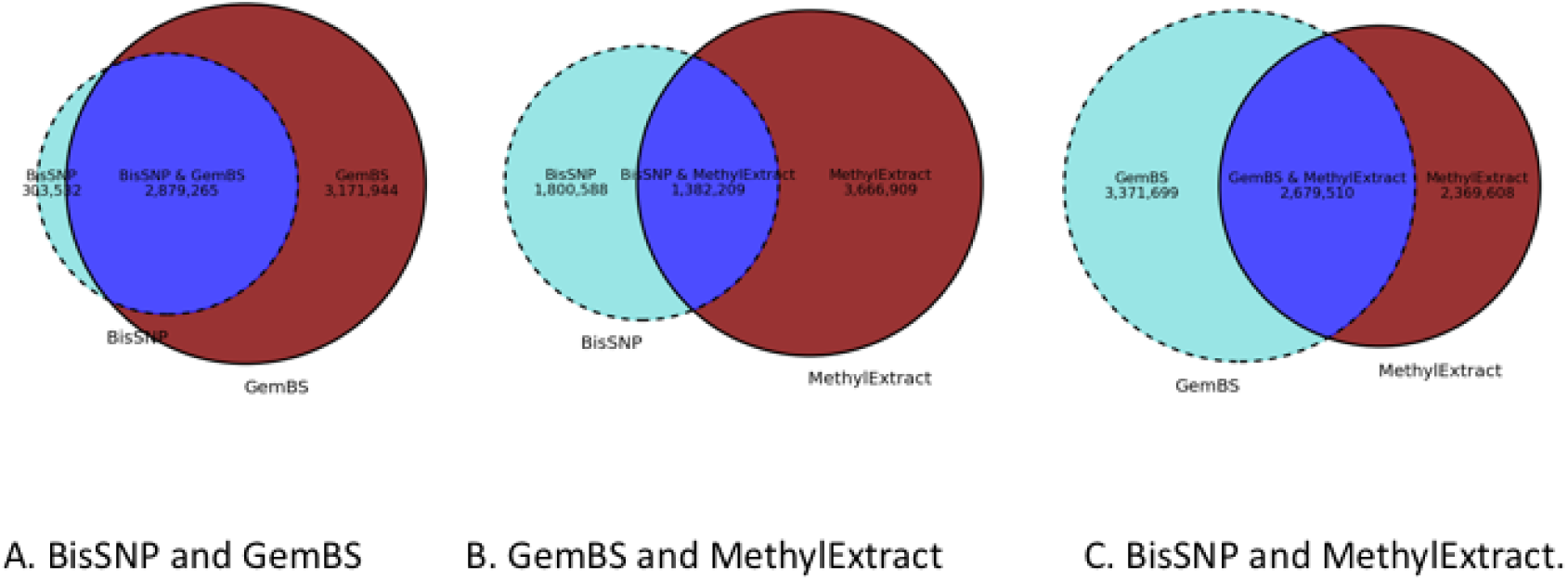
Comparison SNPs based on GEM3 alignments (58X)

**Figure 5.**
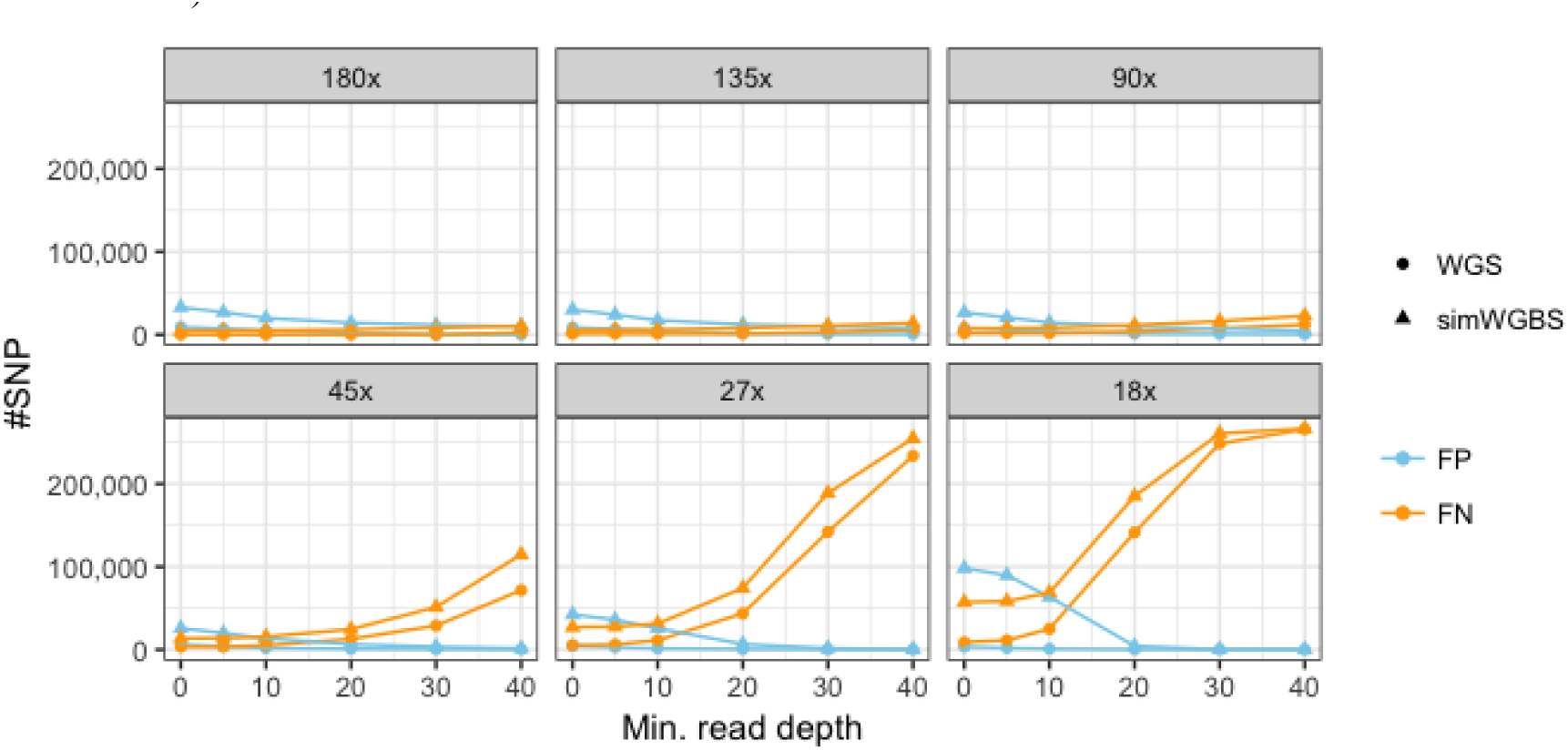
False positive (FP) and false negative (FN) SNP calls from normal sequencing data (WGS) and sequencing data with simulated bisulfite conversion (WGBS) at various sequencing depth and read coverage

In a pairwise comparison, SNP calls generated by BScall and Bis-SNP appear very similar but with a large number of private calls for BScall. Almost all SNPs identified by Bis-SNP are also identified by BScall whereas the overlap with SNPs from MethylExtract is very low (calls based on GEM3 alignment, see Figure 6). BScall and Bis-SNP implement a similar Bayesian model for genotype calling (see above), while MethylExtract identifies SNPs independently of the methylation states. The differences are hence easily explained by different theoretical approaches.

When comparing CpG calls across the three callers, we find Bis-SNP, BScall and MethylExtract detect ~25, ~26, and ~30 million CpGs dinucleotides, respectively. Given only approximately 28 million known CpG sites in the human genome, this suggests a large number of false positive CpG calls resulting from a similarly large number of false positives amongst the private SNP calls reported by MethylExtract (Table 2). This is further supported by the observation that those CpG only reported by MethylExtract are mostly unmethylated and hence likely constitute reference thymines falsely called as cytosines.

**Table 2.**
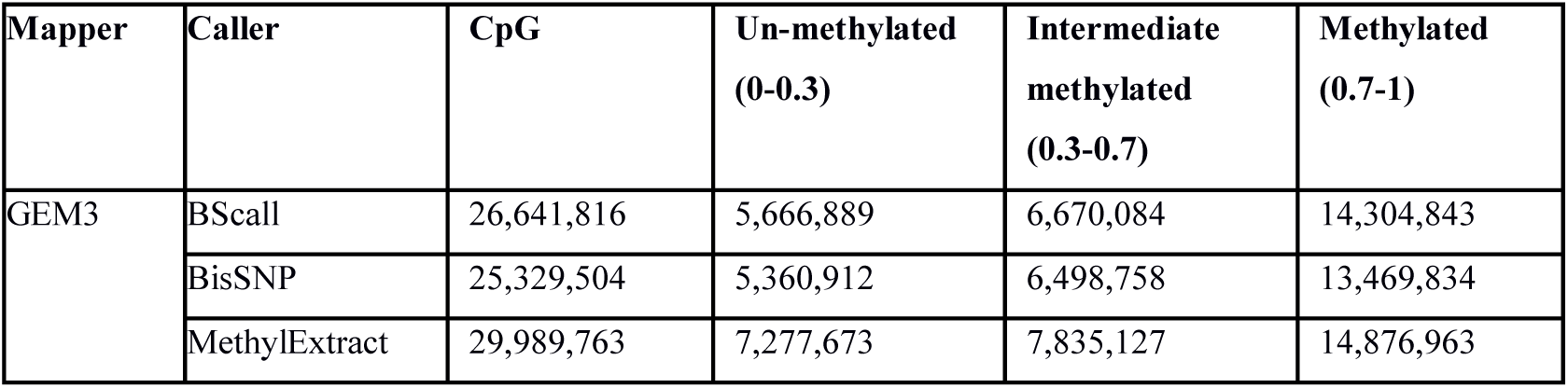
CpG methylation estimates by caller (GEM3 alignment, 58X coverage)

Nevertheless, the vast majority (>85%) of CpG shared across all callers have very similar methylation estimates. This indicates that consistent genotype calls result in consistent methylation estimates. Private CpG calls not shared between BScall and Bis-SNP follow the same distribution of methylation estimates supporting those to be genuine CpG sites. The observed distribution of methylation estimates is expected for this cell type.

Differences in the CpG sites reported by the callers (i.e. private CpGs which are unique to each caller) are difficult to trace. Some might arise through the alignments used, the SNPs identified or simply the way CpG are noted (Table 3.1,3.2). For example, we find that a proportion of the private MethylExtract CpGs are supported by ambiguously mapping reads; while GEMBS only uses uniquely mapped reads. Others might be a result of the larger number of SNPs initially called, such as in the comparison with Bis-SNP and BScall. At the same time SNP calls by Bis-SNP are likely more specific due to the local realignment procedure. Bis-SNP also reports all reference CpG positions whether methylation estimates are available or not (i.e. the position is covered by sufficient uniquely mapped reads or not) leading to an inflated CpG count.

**Table 3.1.** CpG comparision Bis-SNP vs BScall (GEM3 alignment, 58X)

|  | **Total** | **Un-methylated** | **Intermediate methylated** | **Methylated** |
| --- | --- | --- | --- | --- |
| **Shared Equal Methylated** | 23.888.808<br>(85,76%) | 5.093.653<br>(21,32%) | 5.841.360<br>(24,45%) | 12.953.795<br>(54,22%) |
| **Shared Different Methylated** | 853.908<br>(3,06%) |  |  |  |
| **Private BisSNP** | 1.212.855<br>(4,35%) | 286.807<br>(23,64%) | 326.378<br>(26,9%) | 599.670<br>(49,44%) |
| **Private Bscall** | 1.899.100<br>(6,81%) | 415.006<br>(21,85%) | 471.822<br>(24,84%) | 1.012.272<br>(53,30%) |

**Table 3.2.** CpG comparison MethylExtract vs BScall (GEM3 alignment, 58X)

|  | **Total** | **Unmethylated** | **Intermediate methylated** | **Methylated** |
| --- | --- | --- | --- | --- |
| **Shared Equal Methylated** | 26.394.662<br>(87,76 %) | 5.684.621<br>(21,53 %) | 6.853.342<br>(25,96 %) | 13.856.699<br>(52,49 %) |
| **Shared Different Methylated** | 162.186<br>(0,53 %) | - | - | - |
| **Private MethylExtract** | 3.432.915<br>(11,41 %) | 1.573.343<br>(45,83 %) | 860.520<br>(25,06 %) | 999.052<br>(29,02 %) |
| **Private To GemBS** | 84.968<br>(0,28 %) | 66.618<br>(78,40 %) | 10.650<br>(12,53 %) | 7.700<br>(9,06 %) |

### SNP calling from WGBS data

We further tested the utility of BScall for SNP calling from WGBS and the influence of sequencing coverage on the SNP called. We compared SNPs called from whole genome sequencing (WGS) data (using FreeBayes) with SNP calls from simulated WGBS based on the same data set (see METHODS). To cover a wide range of sequence depths and avoid sequencing biases we used data generated for a previous benchmark study (Tyler et al. 2015), namely a medulla sample with ultra-deep sequencing coverage of 180X, which we down sampled to 135x, 90x, 45x, 27x and 18x coverage. BS conversion was simulated taking sequencing and conversion errors into account as well an empirical distribution of methylation observed in real WGBS data of Chronic Lymphatic Leukemia (for details see Methods).

To evaluate the specificity and sensitivity of BScall, we used SNPs called with high confidence from whole genome sequencing (WGS) at maximum sequencing depth (180X, min 1 – 20 reads) as baseline. At this coverage, we find that BScall calls 97% of these SNPs (= true positives) from the simulated WGBS data. Not surprisingly, decreasing coverage results in a smaller number of SNPs being called, both for those called from WGS data and those called from WGBS data (Figure 4). Similarly, the number of false positive SNP call decreases with an increasing read coverage per site, while the number of false negatives increases (Figure 5). The highest number of false SNP calls and missed SNP calls (i.e. FP and FN) are found at the lowest sequencing coverage (18x).

We calculated FDR for all combinations (see SupplTable 5) and determined that, for example, at for WGBS data at a standard sequencing coverage of 27x, a min of 15 reads per site and a FDR of 6.38% BScall covers 84% of the baseline SNPs; at a coverage of 45x and a FDR<5% this increased to 92% (for 27x and 45x WGS data: 92% SNPs with FDR=0.37% and 96% SNPs with 0.51% FDR, respectively). Finally, we checked for incorrect classifications of the SNPs called by BScall. Most errors within the false positive and false negatives are those bases affected by bisulfite treatment, i.e. A>G, C>T, G>A and T>C substitutions (data not shown).

## CONCLUSIONS

We describe GEMBS, a state-of-the-art analysis pipeline for high-throughput bisulfite sequencing data. Compared to other popular tools GEMBS excels in processing speed without sacrificing accuracy during the analysis. GEMBS achieves this through a high-performance bisulfite mapper (GEM3), an efficient genotype caller (BScall) and their able implementation amongst other components. GEMBS execution in an HPC environment can be additionally automated through the use of a meta data server and a pipeline management system. Finally, we demonstrated that GEMBS can accurately call SNPs from WGBS data.

GEMBS is freely available (http://statgen.cnag.es/GEMBS) and has been already used successfully in a number of studies (Kulis et al (2013, 2015), Queiros et al (2016), Schuyler at al (2016), etc.) and several large consortia (e.g. ICGC, BLUEPRINT, PANCANCER).

## Footnotes

1 Note that (a) both the first and second reads can map to either of the two reference sequences: e.g., read 1 can map either to the C -> T reference on the forward strand, or to the G -> A reference on the negative strand and (b) both members of a read pair must map to the same reference sequence (although to different strands).

3 Note: This is contradictory to previous benchmark for Bwa-meth, Pederson et al 2014)

## MATERIAL and METHODS

### Genotype calling

Technically, the BScall model gives the likelihood of the observed bases conditional on genotype, methylation proportion, base error probabilities, and under/over conversion rates. For each possible genotype in turn, BScall calculates the likelihood while maximizing over the unknown methylation proportion. The result is a likelihood profile from which the most likely genotype is selected and reported along with the corresponding maximum likelihood estimate of the methylation proportion

**Figure S1.**
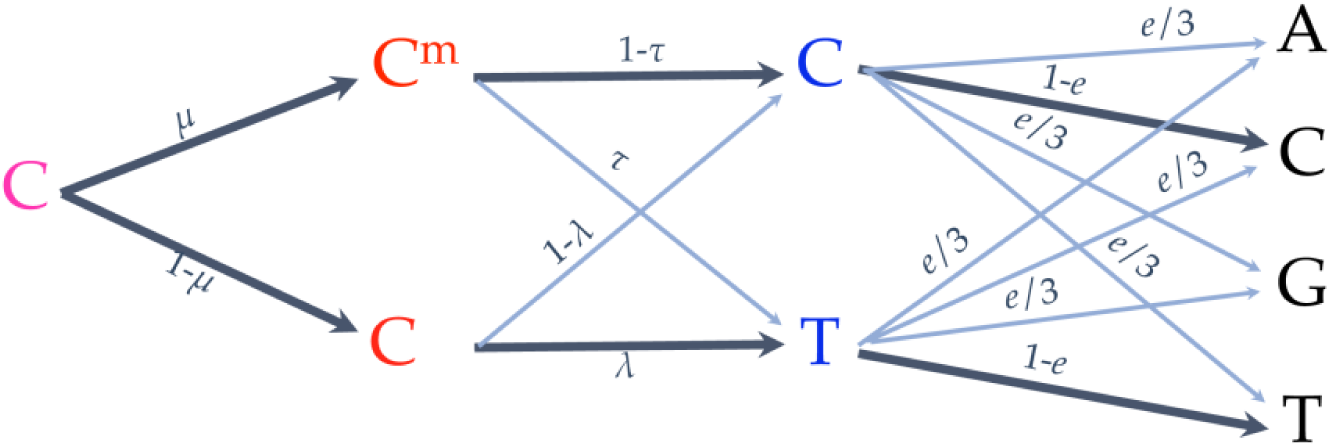
BScall probabilistic model. *P* = likelihood of the observed bases conditional on genotype, methylation proportion, base error probabilities, and under/over conversion rates; *μ*= methylation rate; *λ*= conversion rate; *τ* = over-conversion rate; e = sequencing error probability. Conversion rates for methylated and unmethylated cytosines are determined using spike-in bacteriophage DNA from phage T7 (fully methylated) and phage lambda (unmethylated).

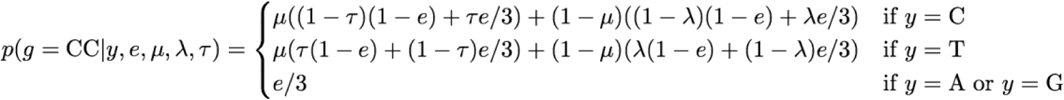

#### Sample data

Read alignments, SNP calls and CpG methylation estimates were benchmarked on WGBS data from a purified human Plasma cell sample extracted from the bone marrow. The sample had been originally sequenced at deep coverage (58X) as part of the European epigenome project BLUEPRINT (www.blueprint-epigenome.eu), but was further down sampled to standard sequencing coverage (27X) for additional comparison.

As baseline data for the simulation (and comparison) we used an ultra-deep sequenced sample of medulla blastoma (180X), which had been used previously in a qualitative control study (Tyler et al. 2012). The high coverage allowed us to test the accuracy of BScall for variant calls at a wide range of coverage depths which we retrieved by down-sampling the original data.

#### Benchmark

All tests have been executed on our in house computing cluster, on nodes of 2 x Xeon E5-2680v3 (12cores each) with 2.5 GHz and 256 GB of main memory using a Linux operative system (Red Hat 6.7).

Alignment software was run in default mode and where available, parameters were used as recommended by the authors (Table 1). Mapping was performed against human genome assembly GRCh38.

**Table 1.**
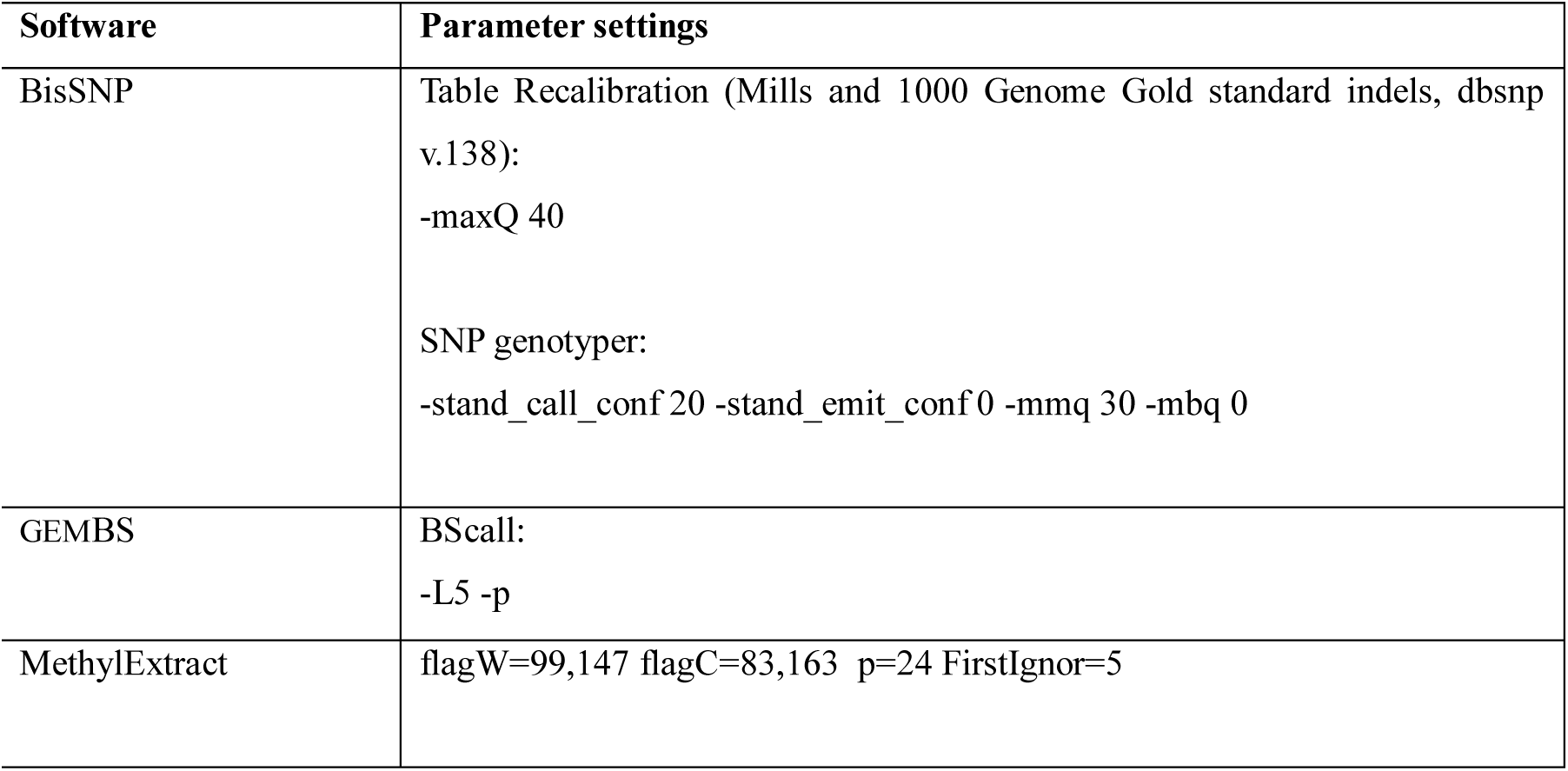
Parameter settings used for alignment

Variant calling software was run in default mode (see Table 2). Each method included marking duplicates prior to the calling step and filtering for CpG positions afterwards. GEMBS (BScall) was executed in parallel by chromosome, Bis-SNP in regions of 10MBp; the results were merged afterwards. For Bis-SNP, we additionally performed local realignment and base quality recalibration as recommend by the authors.

**Table 2.**
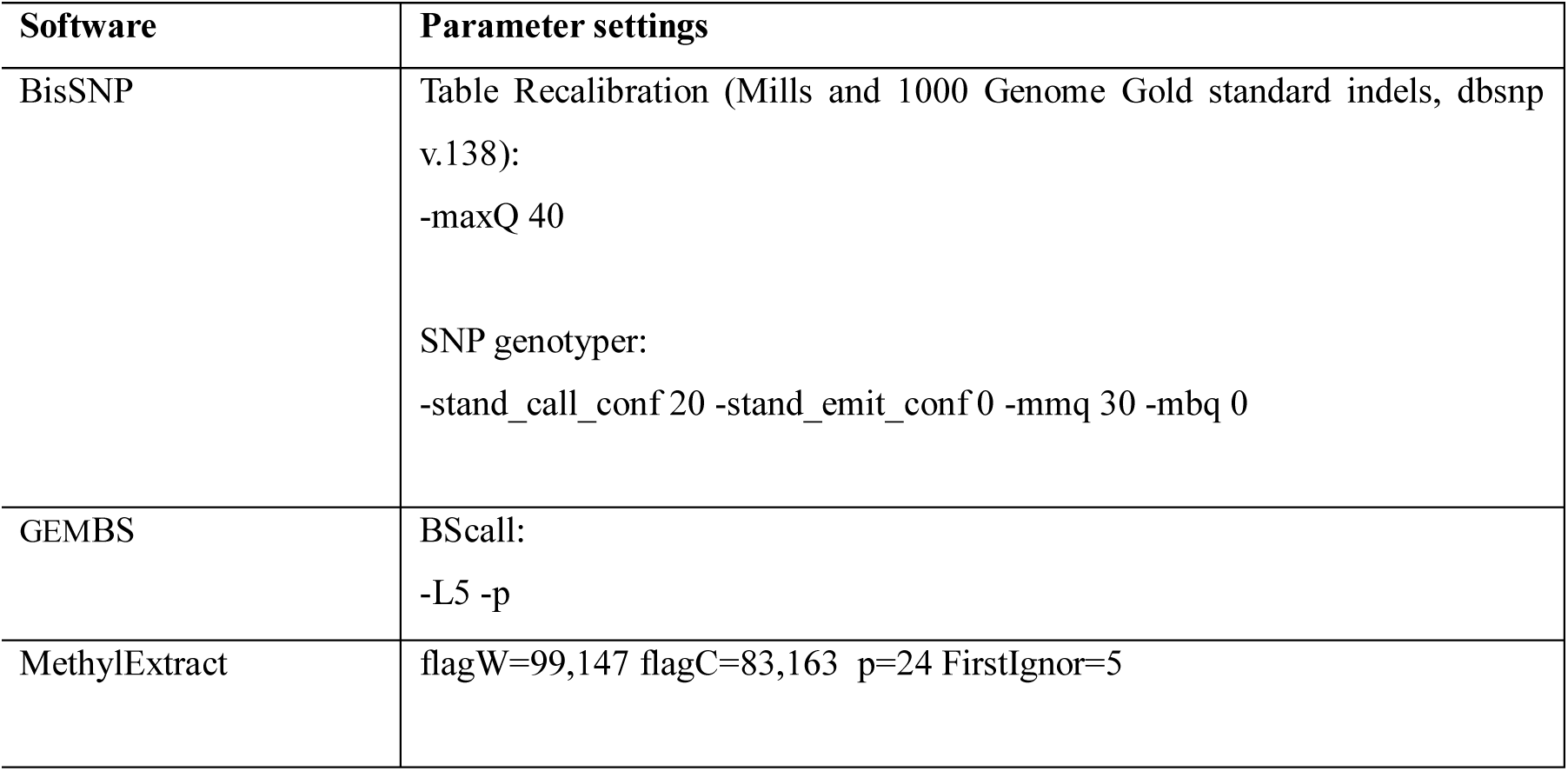
Software and parameter settings used in the variant calling benchmark

#### Variant calling from WGBS

To evaluate the accuracy of BScall to call variants from WGBS data, we compared SNP calls from DNA sequencing data without bisulfite treatments (using FreeBayes) with SNPs called from DNA sequences with simulated bisulfite treatment conversion.

The model we implemented to simulate cytosine conversion takes into account the probability of sequencing errors (e), the probability of the ith cytosine being methylated (m_i_) and the conversion rates for unmethylated cytosines (λ) and methylated cytosines (τ)(see Figure below).

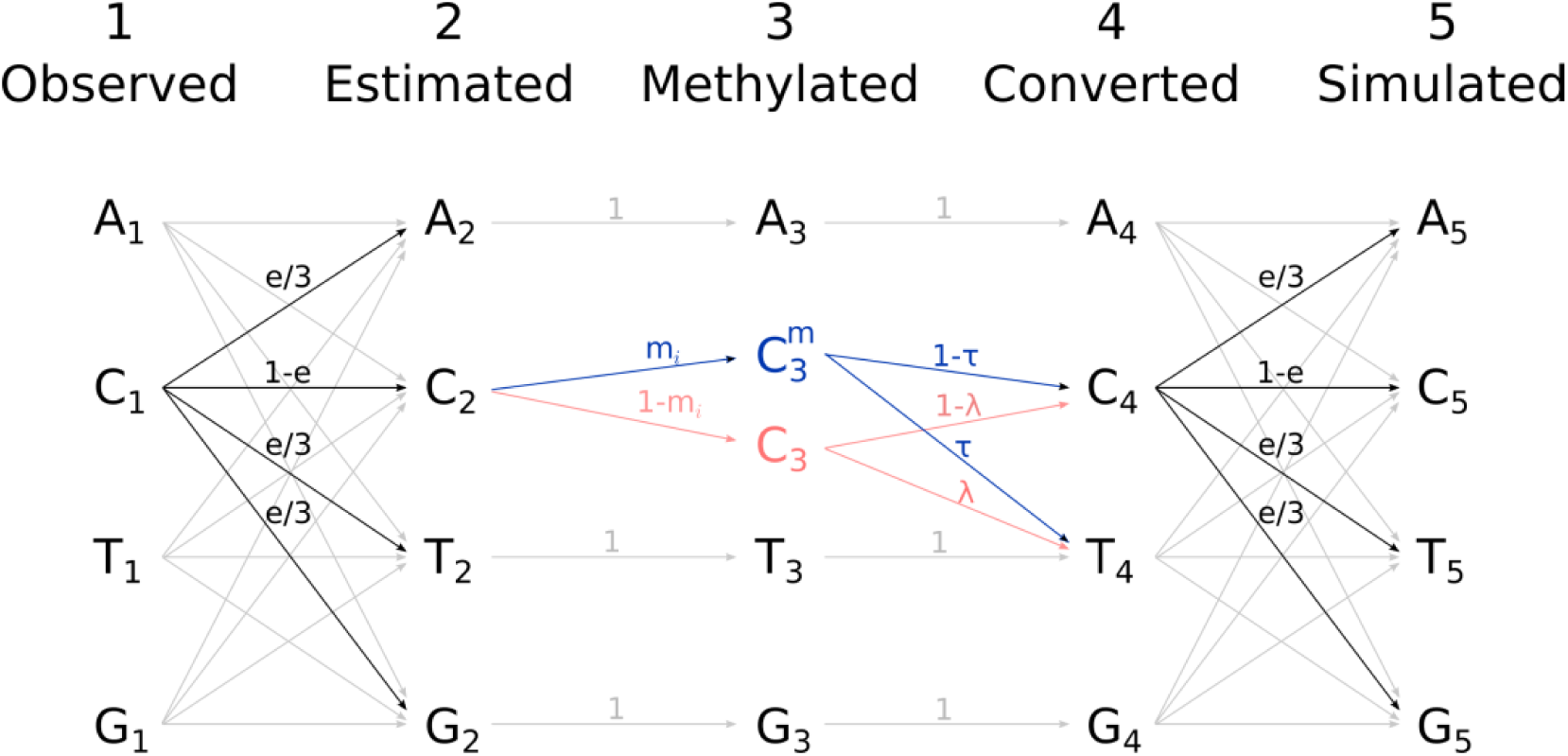

We estimated mi from the methylation profile (= read data) of an available WGBS samples of chronic lymphocytic leukemia (CLL). Missing positions were assigned mi = 0.8 for homozygous CpG, mi = 0.4 for heterozygous CpG and mi = 0 for the remaining genomic positions (assuming that 80% of CpG in the human genome are methylated). Lambda was fixed at λ =0.997 and tau at τ = 0.05 based on our experience. Finally, due to sequencing errors (e) an observed cytosine (C_1_) has a 1-e probability of being a real cytosine (C_2_), and the probability of e/3 of being any other base (A_2_,G_2_,T_2_), all of which need to be taken into account for the simulation.

We can estimate the probability of a position being a particular base by adding the probabilities coming from all the observed bases:

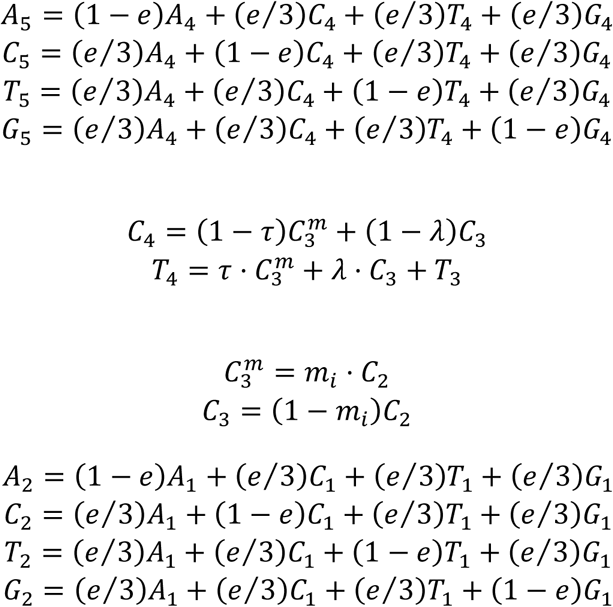

The model has been programmed in C language and is available at https://github.com/bc500/bssim.

## SUPPLEMENT

**Table S1.** Processing time read alignments per mapper for 27X and 58X coverage

| Coverage | Aligner | Elapsed Time (h) | CPU Time (h) |
| --- | --- | --- | --- |
| 27x | Bismark (bowtie2) | 12.3 | 354.3 |
|  | BSMAP | 3.5 | 136.3 |
|  | BwaMeth (bwa-mem) | 6.8 | 278.7 |
|  | GEMBS (GEM3) | 1.1 | 33.5 |
|  | Novoalign | 33.1 | 2419.6 |
| 58x | Bismark (bowtie2) | 25.2 | 732.6 |
|  | BSMAP | 7.4 | 275.6 |
|  | BwaMeth (bwa-mem) | 18.9 | 612.5 |
|  | GEMBS (GEM3) | 2.5 | 69.2 |
|  | Novoalign | 45.5 | 5050.9 |

**Table S2.** Read alignments per mapper for 27X and 58X coverage

| Coverage | Mapper | Bases aligned | Quality > 20 | Read Pairs Mapped | Read Pairs Uniquely Mapped |
| --- | --- | --- | --- | --- | --- |
| 27X | Bismark | 80% | 77% | 87% | 80% |
|  | BSMAP | 85% | 85% | 82% | 82% |
|  | bwa-meth | 97% | 81% | 97% | 84% |
|  | GEM3 | 97% | 85% | 93% | 84% |
|  | Novalign | 86% | 85% | 96% | 88% |
| 58X | Bismark | 80% | 77% | 87% | 80% |
|  | BSMAP | 85% | 85% | 82% | 82% |
|  | bwa-meth | 97% | 81% | 97% | 84% |
|  | GEM3 | 97% | 85% | 93% | 84% |
|  | Novalign | 86% | 85% | 96% | 88% |

**Table S3.** Processing time genotype calling for 27X and 58X coverage

| COVERAGE | CALLER | MAPPER | ELAPSED TIME (h) | CPU TIME (h) |
| --- | --- | --- | --- | --- |
| 27X | Bis-SNP | Bismark | 35.3 | 266.8 |
|  |  | BSMAP | 38.9 | 257.3 |
|  |  | GEM3 | 46.1 | 281.0 |
|  | Bscall | Bismark | 2.2 | 6.3 |
|  |  | BSMAP | 2.0 | 6.2 |
|  |  | GEM3 | 2.7 | 6.4 |
|  | MethylExtract | Bismark | 23.5 | 197.7 |
|  |  | BSMAP | 21.8 | 204.9 |
|  |  | GEM3 | 29.4 | 232.6 |
| 58X | Bis-SNP | Bismark | 65.1 | 419.5 |
|  |  | BSMAP | 60.3 | 405.2 |
|  |  | GEM3 | 92.1 | 471.6 |
|  | Bscall | Bismark | 2.1 | 7.85 |
|  |  | BSMAP | 3.2 | 7.5 |
|  |  | GEM3 | 2.1 | 8.0 |
|  | MethylExtract | Bismark | 45.3 | 357.4 |
|  |  | BSMAP | 42.6 | 372.5 |
|  |  | GEM3 | 55.0 | 421.8 |

**Table S4.** SNP calls per mapper/caller for 27X and 58X coverage

| Coverage | Mapper | Caller | SNPs | Multiallelic<br>SNPs |
| --- | --- | --- | --- | --- |
| 27X | Bismark | Bis-SNP | 2,392,513 | 14,984 |
|  | BSMAP |  | 2,855,296 | 25,914 |
|  | GEM3 |  | 2,716,501 | 23,832 |
|  | Bismark | Bscall | 5,157,821 | 1,577 |
|  | BSMAP |  | 7,284,116 | 6,811 |
|  | GEM3 |  | 5,872,541 | 3,52 |
|  | Bismark | Methyl<br>Extract | 4,093,287 | 3,852 |
|  | BSMAP |  | 4,887,502 | 14,599 |
|  | GEM3 |  | 4,780,534 | 9,282 |
| 58X | Bismark | Bis-SNP | 2,909,705 | 51,76 |
|  | BSMAP |  | 3,448,045 | 72,113 |
|  | GEM3 |  | 3,182,798 | 67,76 |
|  | Bismark | Bscall | 5,138,678 | 1,788 |
|  | BSMAP |  | 7,948,014 | 9,138 |
|  | GEM3 |  | 6,051,209 | 4,22 |
|  | Bismark | Methyl<br>Extract | 4,438,764 | 4,136 |
|  | BSMAP |  | 5,714,662 | 18,399 |
|  | GEM3 |  | 5,049,129 | 9,503 |

**Table S5.** SNP detection for WGS and simulated WGBS data at various coverage levels.

| WGS |  |  |  |  |  | Simulated WGBS |  |  |  |  |
| --- | --- | --- | --- | --- | --- | --- | --- | --- | --- | --- |
| Depth | TP | FP | FN | Sens% | FDR% | TP | FP | FN | Sens% | FDR% |
| 18x | coverage |  |  |  |  |  |  |  |  |  |
| 0 | 258387 | 3406 | 8800 | 96.71 | 1.30 | 210016 | 98090 | 57171 | 78.60 | 31.84 |
| 5 | 255891 | 1988 | 11296 | 95.77 | 0.77 | 208810 | 89757 | 58377 | 78.15 | 30.06 |
| 10 | 242139 | 964 | 25048 | 90.63 | 0.40 | 198628 | 63644 | 68559 | 74.34 | 24.27 |
| 15 | 199969 | 488 | 67218 | 74.84 | 0.24 | 155574 | 24826 | 111613 | 58.23 | 13.76 |
| 20 | 125948 | 233 | 141239 | 47.14 | 0.18 | 82081 | 4297 | 185106 | 30.72 | 4.97 |
| 30 | 18621 | 67 | 248566 | 6.97 | 0.36 | 6494 | 268 | 260693 | 2.43 | 3.96 |
| 40 | 1404 | 37 | 265783 | 0.53 | 2.57 | 526 | 107 | 266661 | 0.20 | 16.90 |
| 27x | coverage |  |  |  |  |  |  |  |  |  |
| 0 | 261905 | 4558 | 5282 | 98.02 | 1.71 | 240118 | 42346 | 27069 | 89.87 | 14.99 |
| 5 | 261020 | 2811 | 6167 | 97.69 | 1.07 | 239453 | 36047 | 27734 | 89.62 | 13.08 |
| 10 | 255912 | 1474 | 11275 | 95.78 | 0.57 | 235851 | 25667 | 31336 | 88.27 | 9.81 |
| 15 | 246021 | 922 | 21166 | 92.08 | 0.37 | 224636 | 15303 | 42551 | 84.07 | 6.38 |
| 20 | 223617 | 515 | 43570 | 83.69 | 0.23 | 193013 | 6646 | 74174 | 72.24 | 3.33 |
| 30 | 125138 | 186 | 142049 | 46.84 | 0.15 | 78410 | 1668 | 188777 | 29.35 | 2.08 |
| 40 | 33460 | 75 | 233727 | 12.52 | 0.22 | 12382 | 315 | 254805 | 4.63 | 2.48 |
| 45x | coverage |  |  |  |  |  |  |  |  |  |
| 0 | 263836 | 5980 | 3351 | 98.75 | 2.22 | 253585 | 25683 | 13602 | 94.91 | 9.20 |
| 5 | 263750 | 4099 | 3437 | 98.71 | 1.53 | 253270 | 20039 | 13917 | 94.79 | 7.33 |
| 10 | 262037 | 1912 | 5150 | 98.07 | 0.72 | 251618 | 13466 | 15569 | 94.17 | 5.08 |
| 15 | 258527 | 1314 | 8660 | 96.76 | 0.51 | 248289 | 9440 | 18898 | 92.93 | 3.66 |
| 20 | 254329 | 974 | 12858 | 95.19 | 0.38 | 242641 | 6526 | 24546 | 90.81 | 2.62 |
| 30 | 238246 | 487 | 28941 | 89.17 | 0.20 | 215862 | 3445 | 51325 | 80.79 | 1.57 |
| 40 | 195596 | 237 | 71591 | 73.21 | 0.12 | 152485 | 1499 | 114702 | 57.07 | 0.97 |
| 90x | coverage |  |  |  |  |  |  |  |  |  |
| 0 | 265405 | 7815 | 1782 | 99.33 | 2.86 | 259696 | 26583 | 7491 | 97.20 | 9.29 |
| 5 | 265405 | 6053 | 1782 | 99.33 | 2.23 | 259570 | 20626 | 7617 | 97.15 | 7.36 |
| 10 | 265389 | 3549 | 1798 | 99.33 | 1.32 | 258912 | 14593 | 8275 | 96.90 | 5.34 |
| 15 | 264909 | 1849 | 2278 | 99.15 | 0.69 | 257646 | 11699 | 9541 | 96.43 | 4.34 |
| 20 | 263333 | 1160 | 3854 | 98.56 | 0.44 | 255770 | 9716 | 11417 | 95.73 | 3.66 |
| 30 | 259500 | 813 | 7687 | 97.12 | 0.31 | 251027 | 7099 | 16160 | 93.95 | 2.75 |
| 40 | 255265 | 596 | 11922 | 95.54 | 0.23 | 244488 | 4724 | 22699 | 91.50 | 1.90 |
| 135x | coverage |  |  |  |  |  |  |  |  |  |
| 0 | 266163 | 8513 | 1024 | 99.62 | 3.10 | 261623 | 29951 | 5564 | 97.92 | 10.27 |
| 5 | 266163 | 6693 | 1024 | 99.62 | 2.45 | 261535 | 23940 | 5652 | 97.88 | 8.39 |
| 10 | 266163 | 4642 | 1024 | 99.62 | 1.71 | 261159 | 17274 | 6028 | 97.74 | 6.20 |
| 15 | 266162 | 2979 | 1025 | 99.62 | 1.11 | 260417 | 14268 | 6770 | 97.47 | 5.19 |
| 20 | 266099 | 1597 | 1088 | 99.59 | 0.60 | 259330 | 12237 | 7857 | 97.06 | 4.51 |
| 30 | 263979 | 634 | 3208 | 98.80 | 0.24 | 256504 | 9843 | 10683 | 96.00 | 3.70 |
| 40 | 261314 | 511 | 5873 | 97.80 | 0.20 | 253205 | 8002 | 13982 | 94.77 | 3.06 |
| 180x | coverage |  |  |  |  |  |  |  |  |  |
| 0 | 267175 | 9032 | 12 | 100.00 | 3.27 | 262552 | 32831 | 4635 | 98.27 | 11.11 |
| 5 | 267175 | 7184 | 12 | 100.00 | 2.62 | 262491 | 26982 | 4696 | 98.24 | 9.32 |
| 10 | 267175 | 5014 | 12 | 100.00 | 1.84 | 262224 | 19860 | 4963 | 98.14 | 7.04 |
| 15 | 267175 | 3392 | 12 | 100.00 | 1.25 | 261709 | 16448 | 5478 | 97.95 | 5.91 |
| 20 | 267175 | 2137 | 12 | 100.00 | 0.79 | 261037 | 14203 | 6150 | 97.70 | 5.16 |
| 30 | 267164 | 25 | 23 | 99.99 | 0.01 | 259085 | 11583 | 8102 | 96.97 | 4.28 |
| 40 | 264840 | 10 | 2347 | 99.12 | 0.00 | 256781 | 9877 | 10406 | 96.11 | 3.70 |

TP, true positives; FP, false positives; FN, false negatives; Sens, sensitivity; FDR, false discovery rate; Depth: minimum read depth;

